## Supplementary figures and images for "Completing Linnaeus’s inventory of the Swedish insect fauna: only 5,000 species left?"

### S1 Figure

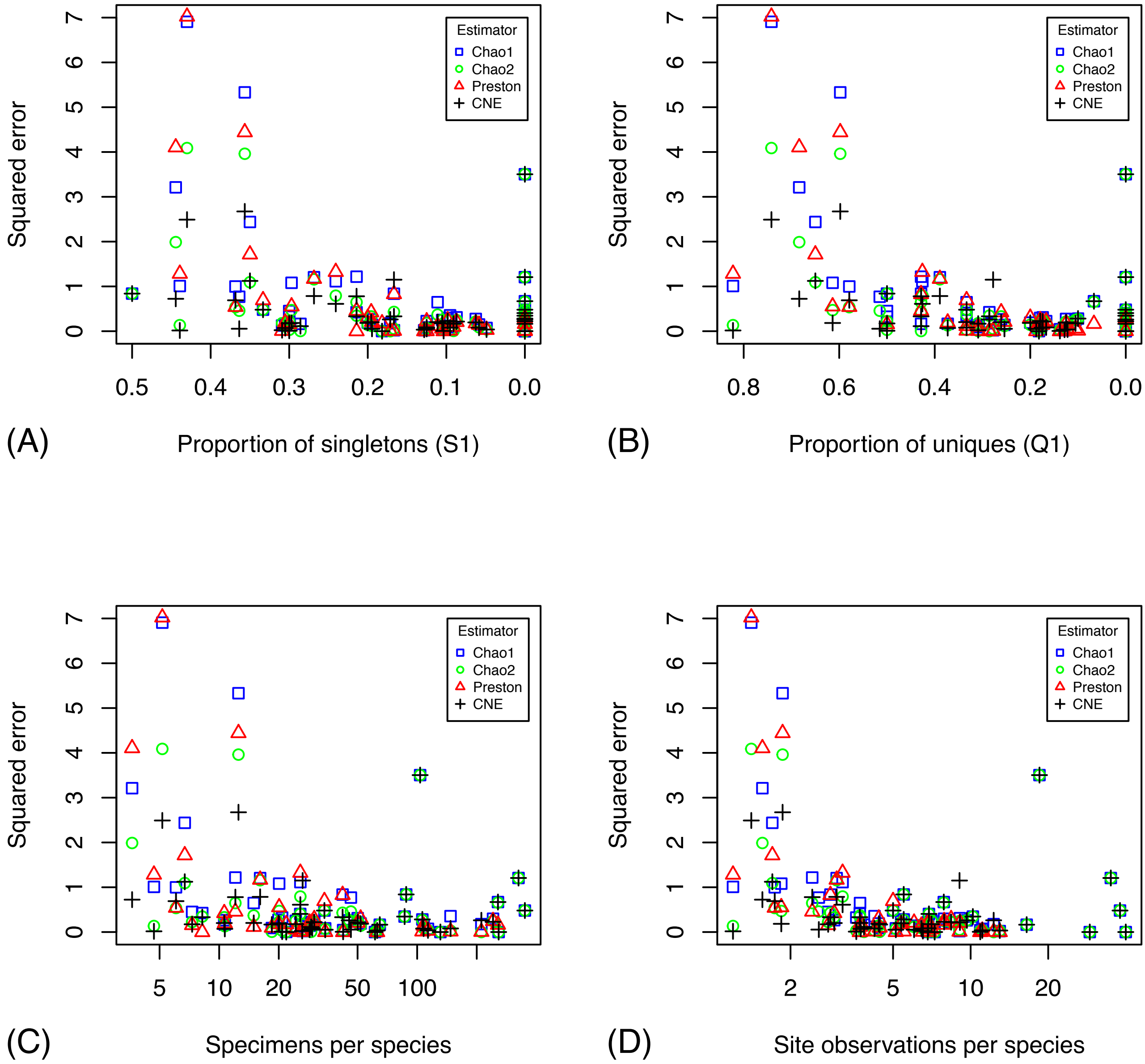

### S2 Figure

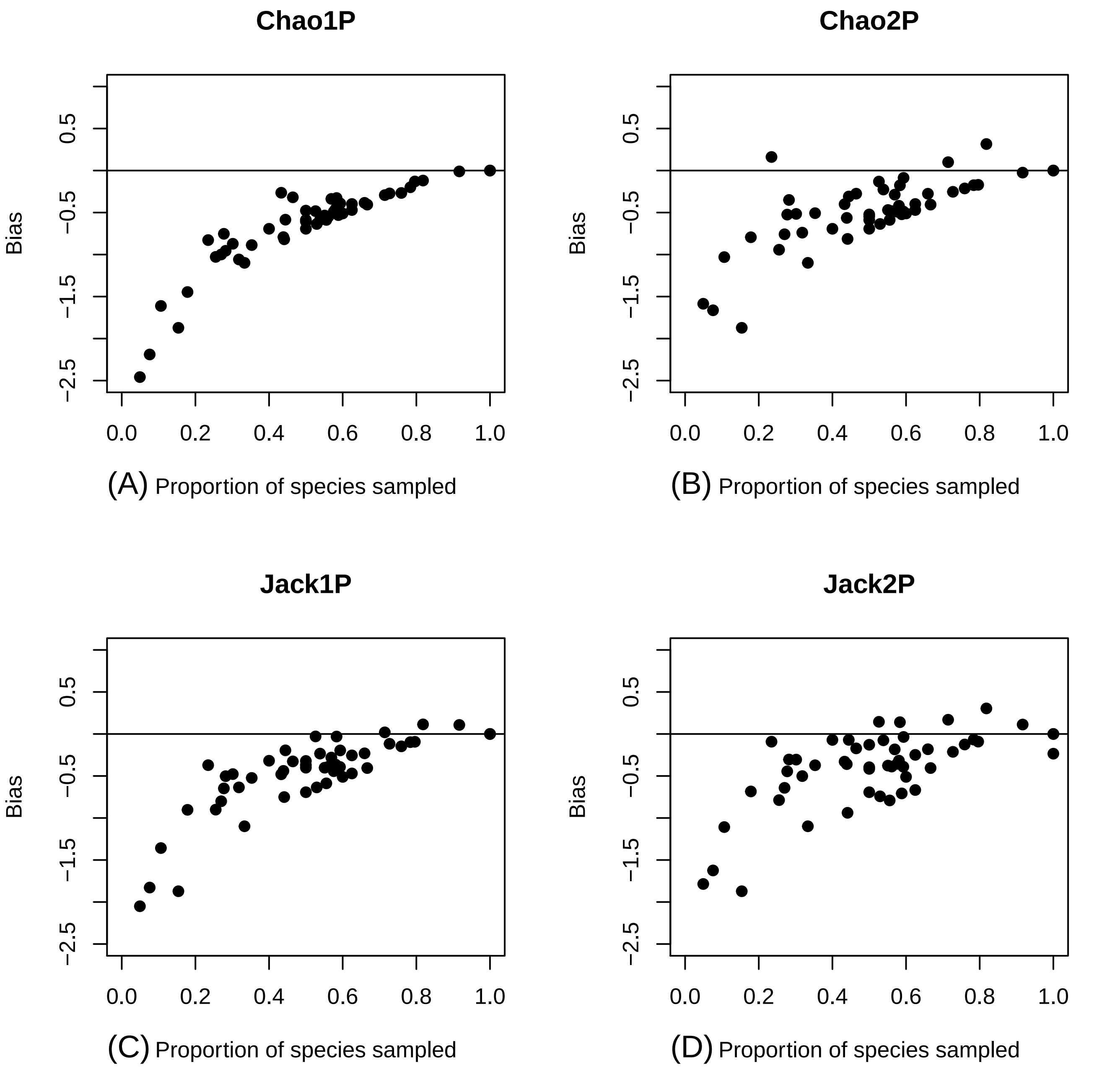
